# Structural Directional Brain–Behavior Asymmetry Across Cortical and Non-Cortical Regions

**DOI:** 10.64898/2026.06.02.729666

**Authors:** Fatemeh Hashemi, Hadis Momtaz, Maryam Malekzadeh, Alireza Kashani

## Abstract

A substantial body of research suggests that the left and right cerebral hemispheres play differential roles in shaping human behavior. However, due to methodological considerations, most studies in this field have relied primarily on psychological methods. Here, we present exploratory neurobiological evidence suggesting that brain volume across numerous cortical and non-cortical homotopic regions may display consistent directional asymmetry in relation to a wide range of behavioral measures. Of particular note, such asymmetry recurred between contralateral homotopic areas across many behavioral parameters. The asymmetric behavioral directionality is distributed across most regions of the human brain, especially in the frontal and temporal cortices, which are particularly developed in humans. This may add a new dimension to previously described aspects of hemispheric asymmetry. In addition, it may help to understand why a marked functional asymmetry can be observed even when structural asymmetry is subtle. Moreover, these findings may shed light on how the brain modulates behavior in health and disease and may contribute to understanding processes involved in neurological and psychiatric disorders.

## INTRODUCTION

Human survival depends on sustaining goal-relevant neural activity, which underlies attention, cognitive engagement, problem-solving, and adaptive decision-making, while suppressing irrelevant stimuli (Zanto & Gazzaley, 2009). In addition, transitions between tasks or mental sets are foundational to organized cognition, enabling flexibility and goal-directed adaptability (Diamond, 2013). This relies on neural mechanisms that allow constant monitoring, modulation, and optimization of cognitive and emotional networks to produce adaptive motor outputs. This homeostatic system, which is crucial for adapting to an ever-changing environment, is underpinned by a delicate balance between excitatory and inhibitory (E/I) neural processes (Dehghani et al., 2016; Knight et al., 1999), yet the precise anatomical bases governing the dynamic bidirectional modulation of neural activity are not fully delineated (Sohal & Rubenstein, 2019; Tatti et al., 2017). The human brain is an asymmetric bi-hemispheric structure. Growing evidence suggests that the left and right hemispheres may differentially influence various aspects of human behavior (Braun et al., 2003; Comer et al., 2015). However, it is not yet clearly understood how bi-hemispheric behavior modulation is carried out. Different studies have proposed a range of mechanisms, from hemispheric specialization and cooperation to competition and opposition (Gazzaniga et al., 1962; Marinsek et al., 2014; Vallortigara & Rogers, 2020). Nevertheless, these mechanisms remain far from conclusive. Thus far, studies on hemispheric modulation of behavior have been mostly based on classic neuropsychological methods, and reports obtained with neurobiological approaches remain comparatively limited (Hervé et al., 2013; Toga & Thompson, 2003). Recent advances in imaging methods have provided detailed information about structural properties of the human brain (Glasser et al., 2016). This provides an opportunity to examine whether hemispheric anatomical characteristics are associated with various behavioral traits. In this study, we used high-resolution structural MRI data from the Human Connectome Project (HCP) to investigate how hemispheric brain volume in cortical and non-cortical regions correlates with various behavioral measures.

## MATERIALS AND METHODS

All structural and behavioral measures were obtained from the HCP 1200-subject release, as described previously (Malekzadeh & Kashani, 2020; Van Essen et al., 2013). Structural MRI brain data and behavioral measures from 1,113 participants, including 606 females and 507 males aged 22–35 years, were analyzed (Source Data 1). Statistical analyses were performed using SPSS software version 24.0 (IBM, Armonk, NY, USA). We initially considered 34 bilateral cortical gray-matter volumes according to the Desikan–Killiany atlas, seven bilateral non-cortical volume pairs, cerebellar gray and white matter volumes, total intracranial volume, age, and gender as potential predictors. Data were checked for linearity, normality of the residuals, homoscedasticity, and multicollinearity to verify the assumptions of multiple linear regression (Osborne & Waters, 2002). Variance Inflation Factor (VIF) threshold was set to 10 (Neter et al., 1990).

The caudate nucleus and cerebellar gray matter volume were excluded because of multicollinearity. To comprehensively examine the unique contributions of individual brain regions to the behavioral measures, we employed a simultaneous (forced-entry) multiple linear regression model including all remaining predictors. This approach was deliberately chosen over dimensionality-reduction techniques such as principal component analysis, because such methods, while mitigating multicollinearity, collapse anatomically distinct structures into latent components or summary metrics and thereby reduce the regional specificity central to interpreting structure–function relationships in the human brain (Mwangi et al., 2014). Although inclusion of all predictors may increase multicollinearity, yield modest overall explanatory power, and reduce the stability of individual regression coefficients, this interdependence likely reflects the biological covariance structure of brain morphometry rather than merely a removable statistical nuisance (Gregorich et al., 2021; Smith & Nichols, 2018). We therefore retained the full predictor set to estimate each region’s partial association while controlling for the others. This model was used to explore the relationship between brain structure and each of the 147 behavioral measures treated as dependent variables. To assess the robustness of the associations and account for potential instability due to multicollinearity, we also calculated partial, part, and zero-order coefficients. The statistical significance level was set at p < 0.05. This was an exploratory analysis; therefore, p-values were not corrected for multiple comparisons (Bender & Lange, 2001). The statistically significant hemispheric standardized regression β coefficients across all models were considered for further analysis. Hereafter, unless otherwise specified, the term “brain region(s)” refers to the regional volumes specified above.

## RESULTS

The models examined in this study revealed a range of outcomes, consistent with typical findings in neuroimaging research on brain morphometry and behavioral measures, spanning from minimal explanatory power with no statistical significance to significant associations with more substantial explanatory power (see Extended Data 1 for full details and Extended Data 2 for a summary).

Statistically significant predictors across all models included 803 brain-region, 58 gender, 32 age, and 12 intracranial-volume variables. Across all models, 70% of zero-order and 100% of part and partial correlation coefficients exhibited the same direction as the β coefficients.

Of regional β coefficients that were significantly associated with a measure, 80% showed a unilateral distribution, with 49% linked to the left hemisphere and 51% to the right hemisphere. In contrast, 20% displayed a bilateral pattern, i.e., both contralateral homotopic regions were significantly associated with the measure (Extended Data 2).

The directional aspect of the relationships was consistent across similar measures. For example, when significant, the left superior temporal cortex (STG) always correlated negatively with accuracy and positively with reaction time across all cognitive tests (Extended Data 1). We refer to this as Regional Valence (RV). When both contralateral homotopic regions were associated with a measure, directional opposition in RV was observed in almost all cases (97%); i.e., the left and right hemispheres were inversely correlated with the measure. We describe this as directional asymmetry (DA) (Fig. 1–4). For example, RV in the left and right STG consistently opposed each other relative to numerous working memory reaction time measures (WMRT) (Fig. 3). This will hereafter be referred to as homotopic directional asymmetry (hDA).

**Fig. 1A.**
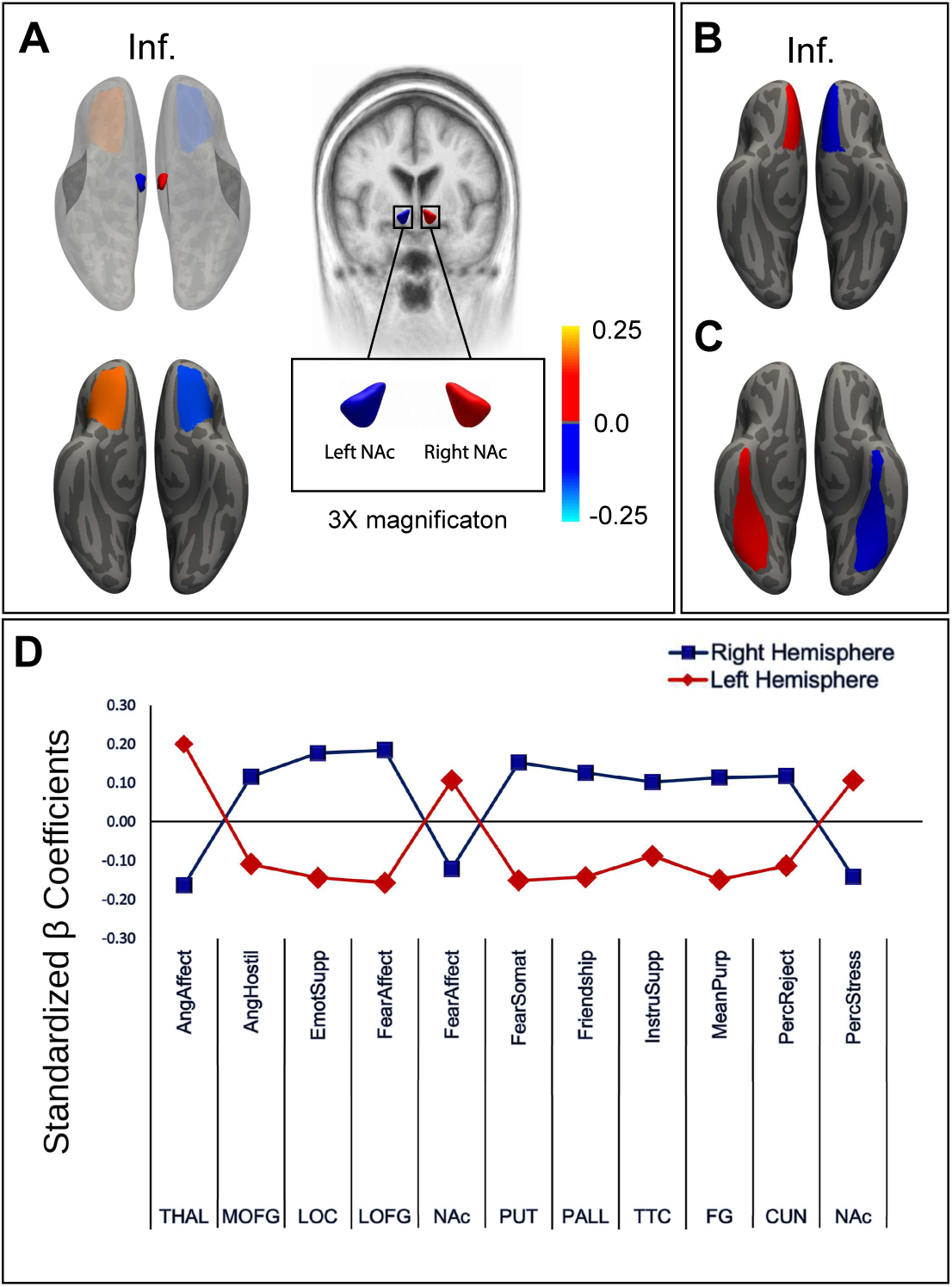
Directional Asymmetry (DA) in Emotion and Psychosocial Measures. Surface- and volume-based maps show standardized β coefficients illustrating: 1. Crossed multiregional DA with dual hDA and iDA components: (A) Lateral orbitofrontal gyrus and nucleus accumbens versus NIH Toolbox Fear-Affect Survey (FearAffect). The brain background was adjusted to enhance the visibility of the nucleus accumbens. 2. Regional DA: (B) Medial orbitofrontal gyrus versus NIH Toolbox Anger-Hostility Survey (AngHostility). (C) Fusiform gyrus versus NIH Toolbox Meaning and Purpose Survey (MeanPurp). (D) Scatter plot showing standardized β coefficients exhibiting DA across various brain regions relative to emotion and psychosocial measures.

**Fig. 1B.**
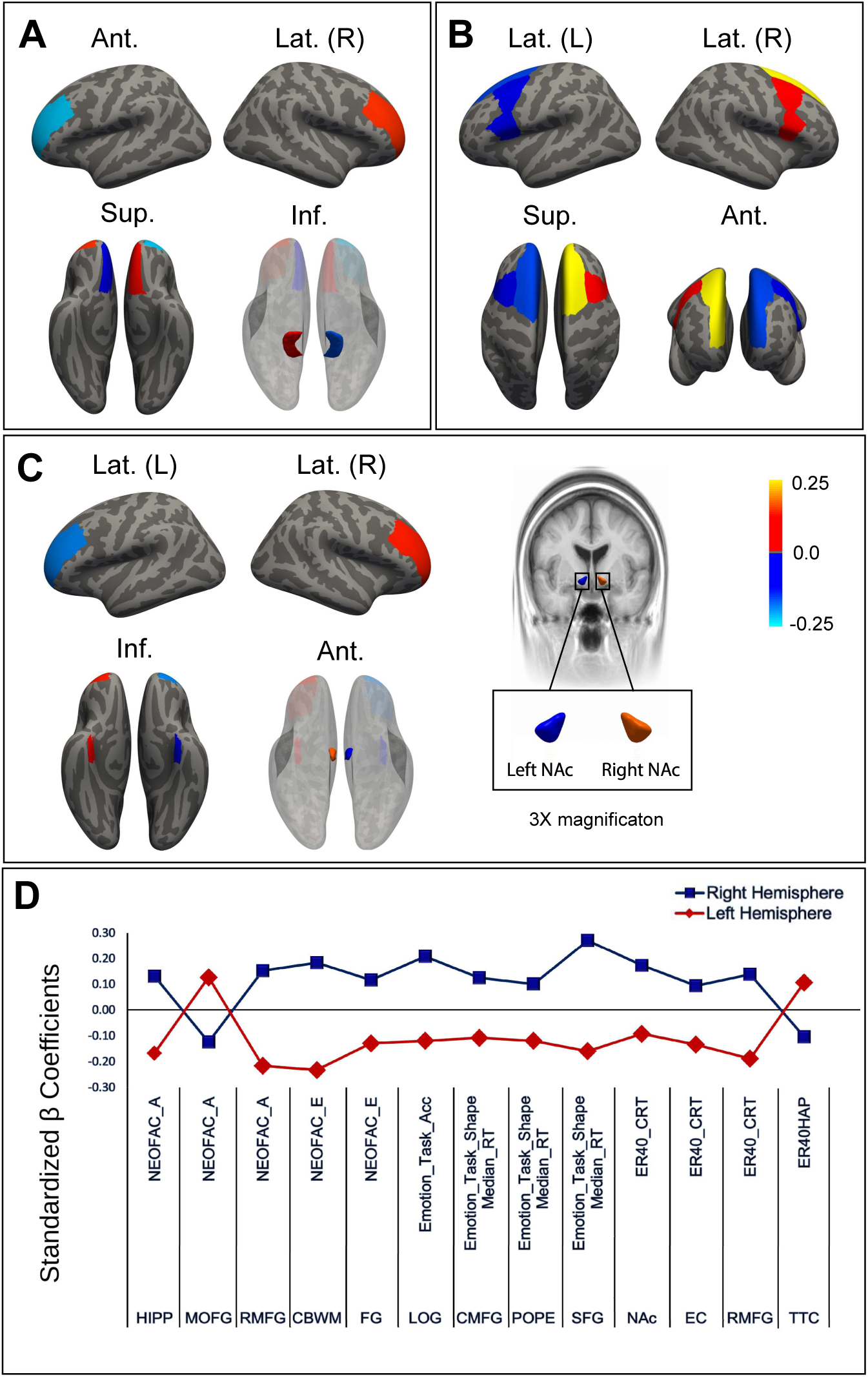
DA in Personality and Emotion Task Measures. Surface- and volume-based maps show standardized β coefficients illustrating: 1. Crossed multiregional DA with dual hDA and iDA components: (A) Rostral middle frontal and medial orbitofrontal gyri and hippocampus versus NEO Five-Factor Inventory Agreeableness (NEOFAC_A). 2. Concordant multiregional DA with dual hDA and iDS components: (B) Superior frontal and caudal middle frontal gyri versus Emotion Recognition Task Shape Median Reaction Time. (C) Rostral middle frontal and entorhinal gyri and nucleus accumbens versus Penn Emotion Recognition 40 (ER-40) Correct Response Time (CRT). The brain background was adjusted to enhance the visibility of the nucleus accumbens. (D) Scatter plot showing standardized β coefficients exhibiting DA across various brain regions relative to personality and emotion task measures.

**Fig. 2.**
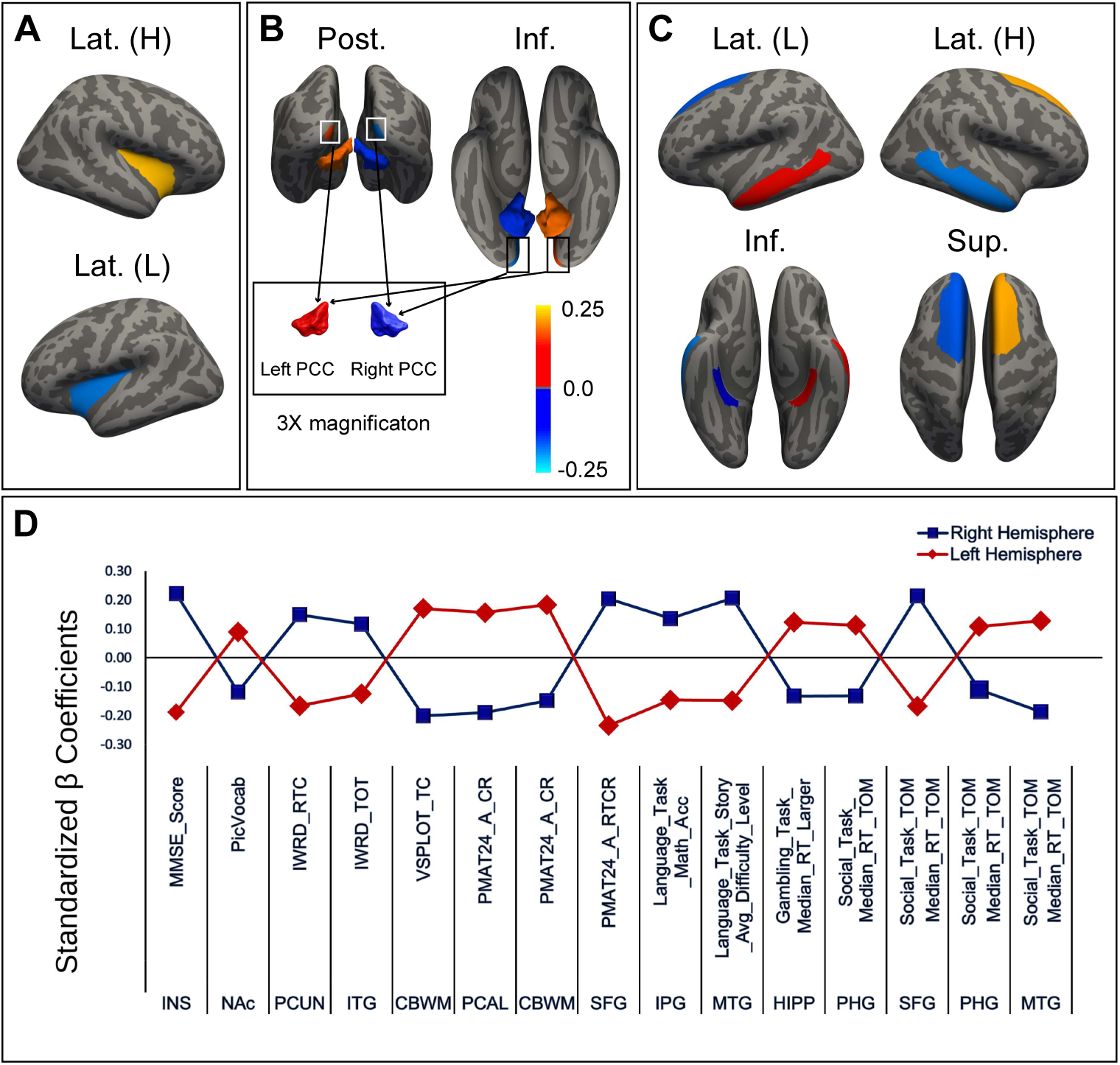
DA in Cognitive and Task Performance Measures. Surface- and volume-based maps show standardized β coefficients illustrating: 1. Regional DA: (A) Insula versus Mini-Mental State Examination (MMSE) Score. 2. Concordant multiregional DA with dual hDA and iDS components: (B) Cerebellar white matter and pericalcarine cortex versus Penn Matrix Reasoning Test 24-item Accuracy (PMAT24-A_CR). 3. Crossed multiregional DA with dual hDA and iDA components: (C) Middle temporal, superior frontal, and parahippocampal gyri versus Social Task Theory of Mind Median Reaction Time (Social_Task_TOM_Median_RT_TOM). (D) Scatter plot showing standardized β coefficients exhibiting DA across various brain regions relative to cognitive and task performance measures.

**Fig. 3A.**
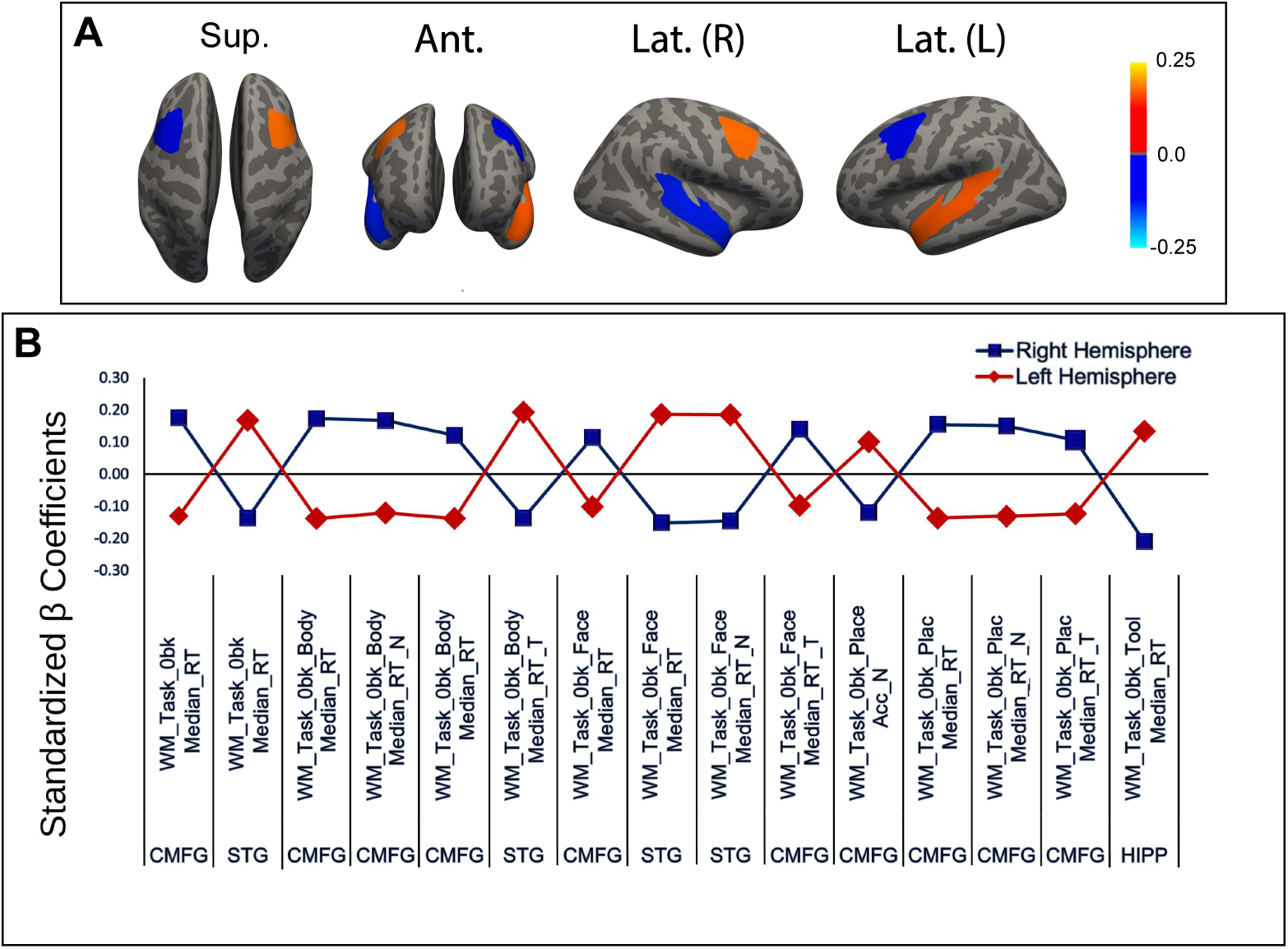
DA in 0-Back Working Memory Reaction Time Measures. Surface- and volume-based maps show standardized β coefficients illustrating: 1. Crossed multiregional DA with dual hDA and iDA components: (A) Caudal middle frontal and superior temporal gyri versus 0-back Working Memory Task Median Reaction Time. (B) Scatter plot showing standardized β coefficients exhibiting DA across various brain regions relative to 0-back Working Memory Task Median Reaction Time Measures.

**Fig. 3B.**
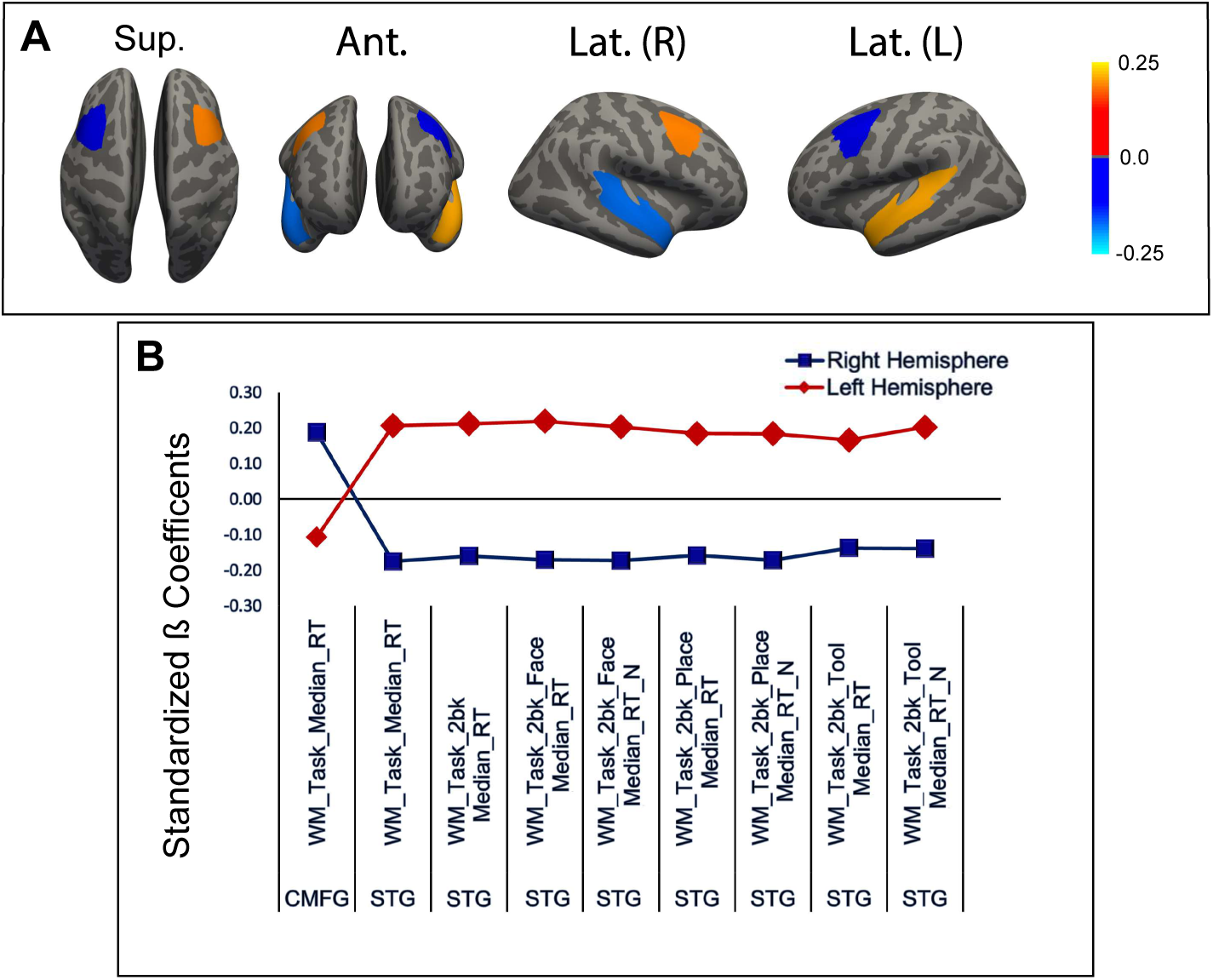
DA in Total and 2-Back Working Memory Reaction Time Measures. Surface- and volume-based maps show standardized β coefficients illustrating crossed multiregional DA with dual hDA and iDA components: (A) Caudal middle frontal and superior temporal gyri versus Total Working Memory Task Median Reaction Time (WM_Task_Median_RT). (B) Scatter plot showing standardized β coefficients exhibiting DA across various brain regions relative to Total and 2-back Working Memory Task Median Reaction Time measures.

**Fig. 3C.**
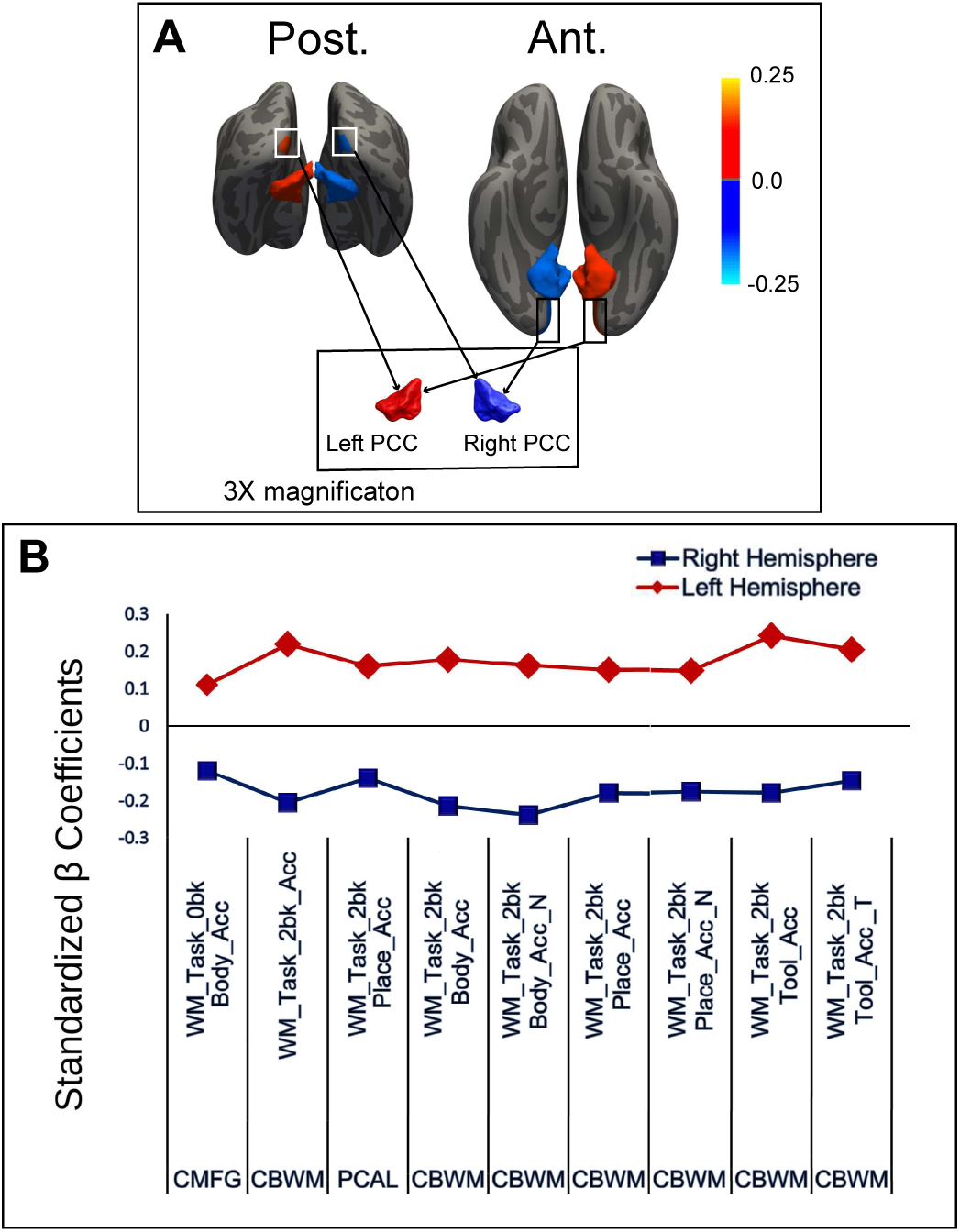
DA in Working Memory Accuracy Measures. Surface- and volume-based maps show standardized β coefficients illustrating concordant multiregional DA with dual hDA and iDS components: (A) Pericalcarine cortex and cerebellar white matter versus Working Memory Task 2-Back Accuracy. (B) Scatter plot showing standardized β coefficients exhibiting DA across various brain regions relative to working memory accuracy measures, including Total 2-Back Accuracy.

**Fig. 4.**
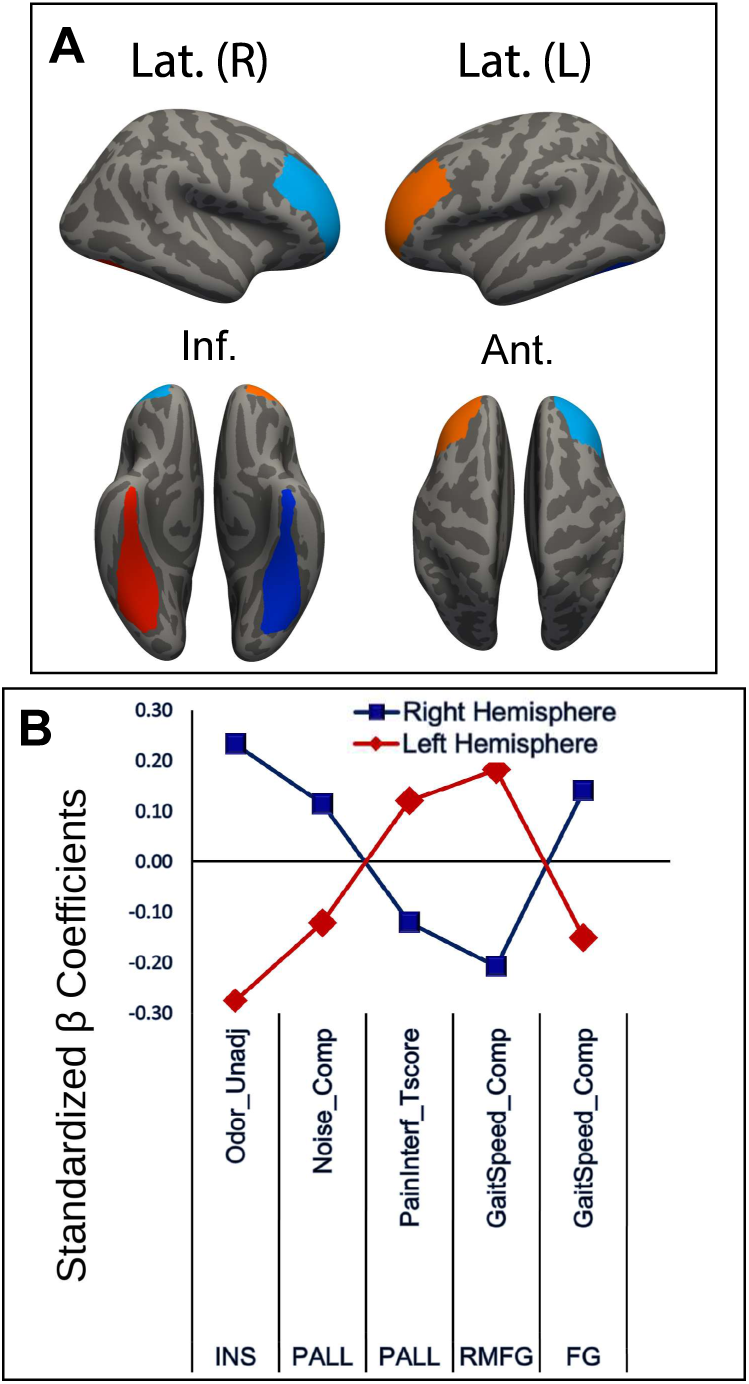
DA in Sensorimotor Measures. Surface- and volume-based maps show standardized β coefficients illustrating crossed multiregional DA with dual hDA and iDA components: (A) Rostral middle frontal and fusiform gyri versus Gait Speed Composite (GaitSpeed Comp). (B) Scatter plot showing standardized β coefficients exhibiting DA across various brain regions relative to sensorimotor measures.

Only two brain regions, the posterior cingulate and entorhinal cortices, presented a concordant homotopic RV, exhibiting directional symmetry (hDS), relative to two outcomes (3%) (Extended Data 2).

hDA was observed in the relationships between 59% of cortical regions (20/34) with at least one outcome. A total of 61 behavioral measures (41%) exhibited associations characterized by hDA (Fig. 1–4). They include 16 emotion and personality, 7 cognition, 28 WM, 7 social and gambling, 4 sensorimotor, and 2 language measures (Extended Data 2, Fig. 1–4). The distribution of the cortical regions presenting hDA is as follows: 40% in the frontal cortex, 42% in the temporal cortex, 4% in the parietal cortex, 9% in the occipital cortex, and 2% in the insula (Fig. 5). All analyzed subcortical regions except amygdala (85%) displayed hDA with at least one measure.

**Fig. 5.**
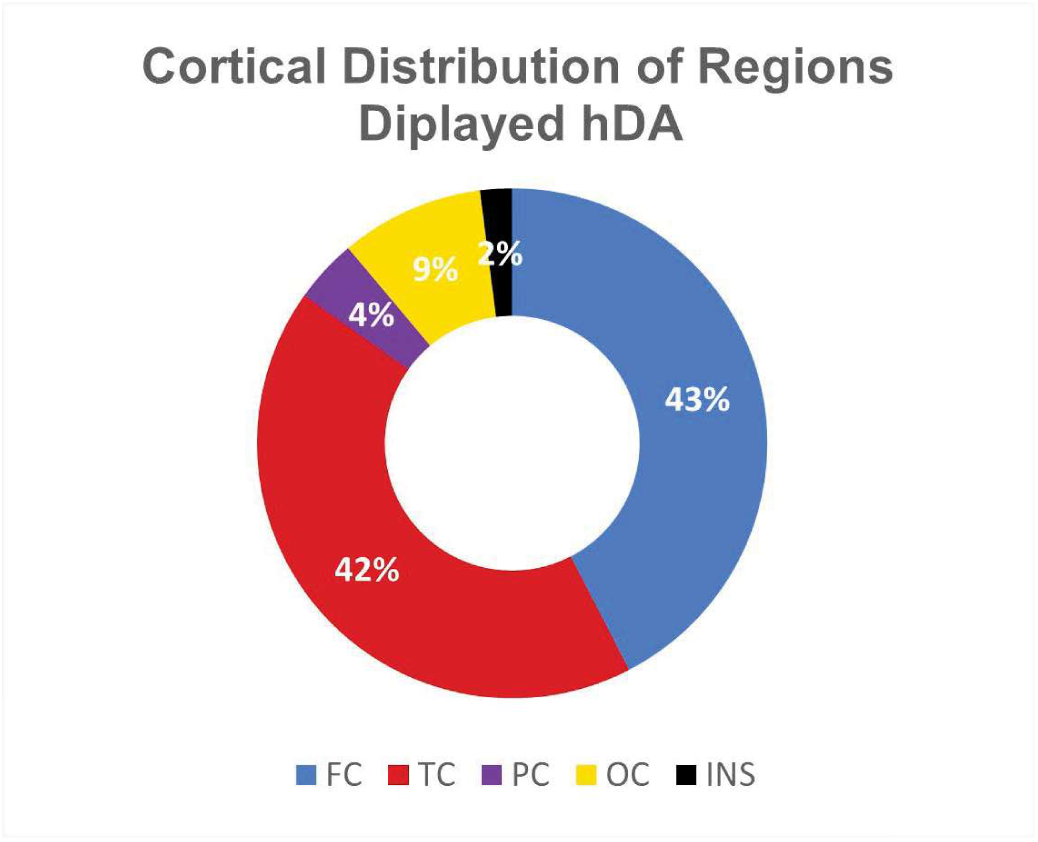
Distribution of the cortical regions that exhibited hDA.

In several cases, multiregional DA patterns were observed, including 9 instances involving bi-regional four-hemisphere and 4 instances involving tri-regional six-hemisphere associations (Fig. 1–4, 8A–C). In all cases, hDA was consistently observed. However, interregional ipsilateral relationships were either crossed or concordant. Specifically, ipsilateral directional asymmetry (iDA) was observed in 67% (6/9) of bi-regional associations. However, in 33% (3/9) the ipsilateral hemispheric relationships were concordant (iDS). The first case creates a crossed multiregional DA with dual hDA and iDA components. In other words, in addition to contralateral homotopic regions, ipsilateral heterotopic areas oppose each other in relation to the measure. For instance, the caudal middle frontal gyrus (cMFG) and STG presented a consistent crossed DA in relation to the total median WMRT measure (Fig. 3B; Fig. 8C). Therefore, consistent asymmetric organization of RV may be observed ipsilaterally as well. In fact, iDA was observed across a broad range of behavioral domains, including emotion, working memory, cognition, and sensorimotor measures (Fig. 1–4). Of tri-regional relationships, in two cases, all ipsilateral hemispheres were exclusively symmetric, exhibiting iDS (Fig. 1B). Conversely, in two instances concomitant iDS and iDA were noticed. For example, the rostral middle frontal (rMFG) and medial orbitofrontal gyri (MOFG) exhibited a crossed DA with respect to NEO Five-Factor Inventory Agreeableness (NEOFAC_A). However, the hippocampus demonstrated a concordant DA with the rMFG and a crossed DA with MOFG relative to the same measure (Fig. 1B, Fig. 8). The same pattern was observed between the middle temporal (MTG), superior frontal (SFG) and parahippocampal gyri in relation to social task theory of mind median reaction time (Fig. 2, Fig. 8B). Notably, in both of these cases of tri-regional associations, regions exhibiting only iDA were located in the neocortex, and the third region exhibiting iDS with one of the neocortical regions was located in the allocortex (Fig. 8A,B).

Regions that exhibited simultaneous iDA and hDA had the following distributions: frontal lobe (56%), temporal lobe (39%), and non-cortical regions (11%) (Fig. 6). The distribution of regions exhibiting concomitant hDA and iDS was: frontal lobe (33%), temporal lobe (17%), occipital lobe (17%), and non-cortical regions (33%) (Fig. 7).

**Fig. 6.**
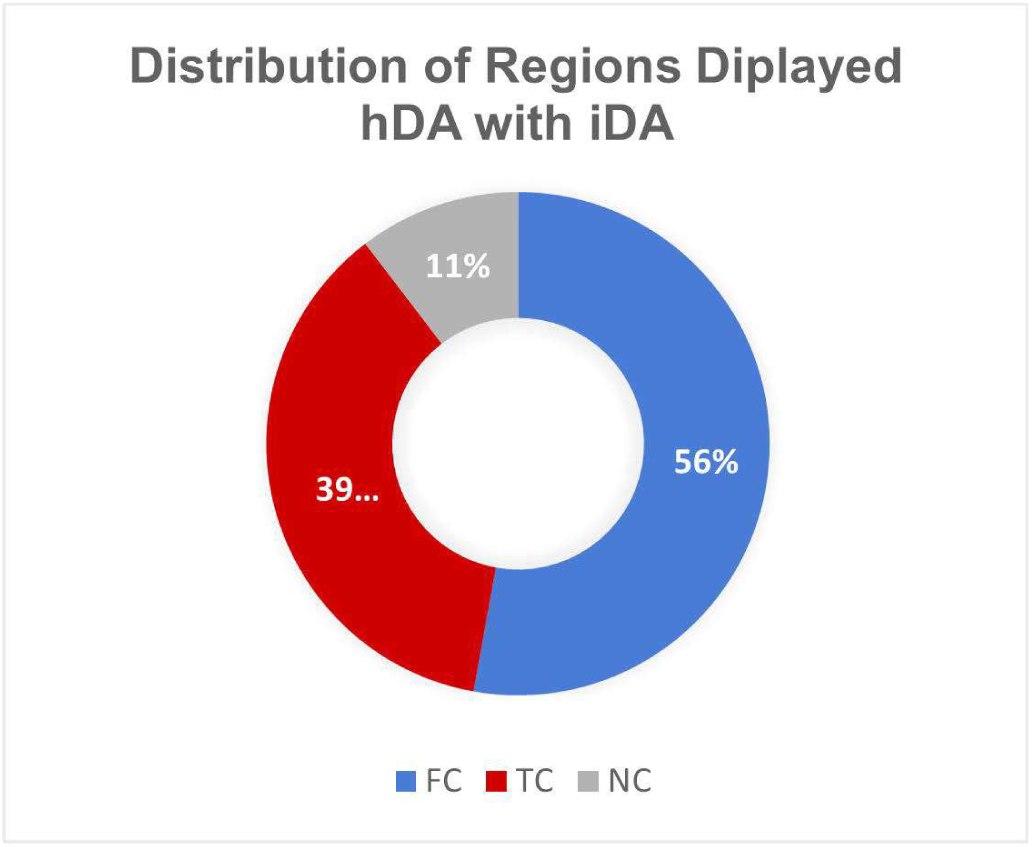
Distribution of the regions that exhibited simultaneous hDA and iDA.

**Fig. 7.**
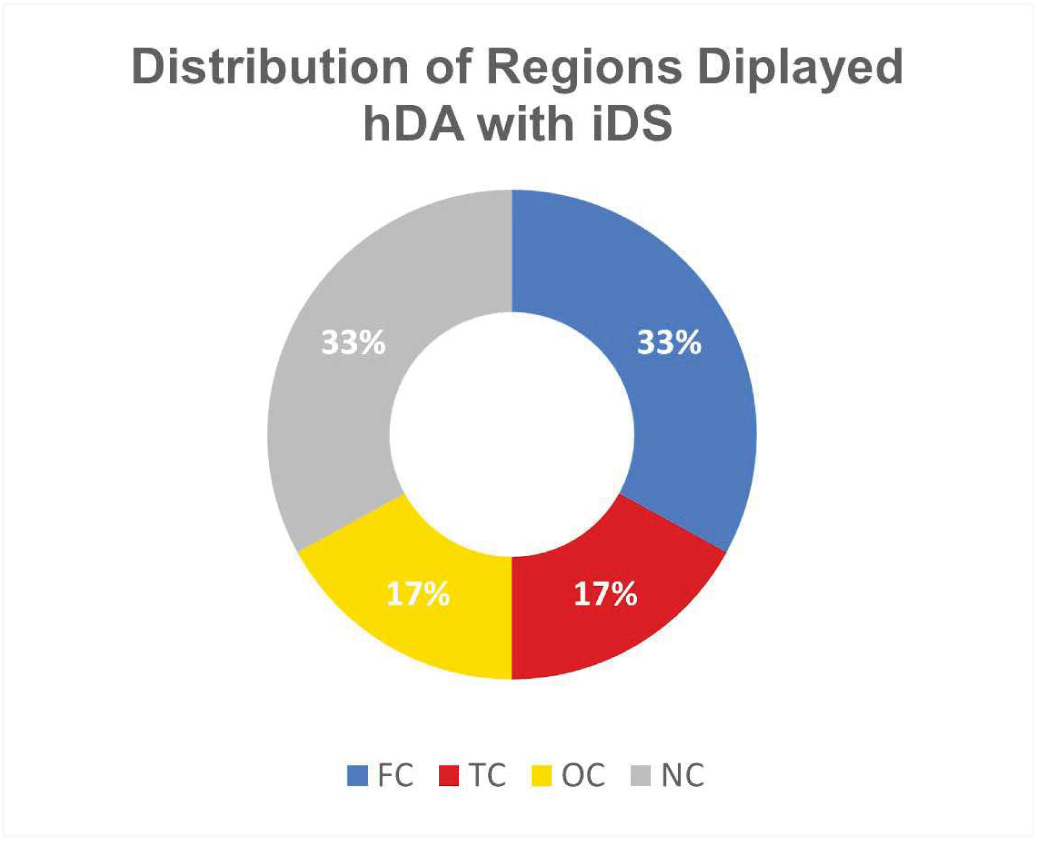
Distribution of the regions that exhibited simultaneous hDA and iDS.

Notably, iDA was not limited to cases involving crossed multiregional DA. Rather, consistent iDA was also observed independently across the same or similar behavioral measures. For instance, the left lingual gyrus consistently opposes the left pericalcarine cortex relative to all cognitive measures when both regions show significant effects (Fig. 9).

## Discussion

The present study examined associations between morphometric features of the left and right cerebral hemispheres and a broad range of behavioral outcomes. In multiple-regression analyses such as those used here, some models may show limited explanatory power and may not reach overall statistical significance. Notably, the omnibus F-test and tests of individual regression coefficients address different hypotheses: the omnibus test evaluates the joint contribution of all predictors, whereas each coefficient test evaluates the unique association of a given predictor after adjustment for the others. Accordingly, a non-significant omnibus model does not necessarily preclude interpretation of individually significant coefficients. This is especially relevant in brain–behavior research, where associations are often small, distributed, and multifactorial. In such settings, individual coefficients may capture localized or theoretically meaningful associations even when the variance explained by the full model is modest. As Gregorich et al. argue, omission of relevant correlated predictors should not be the default response to multicollinearity, because excluding such variables can bias parameter estimates when they are necessary to address the research question (Gregorich et al., 2021). Retaining theoretically motivated predictors may therefore provide a more complete neuroanatomical profile and better align the analysis with a hypothesis-driven framework that prioritizes biological interpretability rather than purely predictive optimization. Within this framework, we believe that interpreting individual coefficients showing significant associations with the measures, together with their consistent patterns across models, may yield meaningful insight into the brain’s role in the fine-tuning of human behavior. For this reason, individually significant predictors were examined even when the overall model F-test did not reach statistical significance. Importantly, for each of the predictors, the direction of the standardized β coefficient was identical to the direction of both its partial and part correlations. Because all these correlations are mathematically related measures of the adjusted effect of a predictor while controlling for all others, the complete directional agreement supports the reliability of the estimated directions (Cohen et al., 2003; Kutner et al., 2005). In addition, the fact that approximately 70% of zero-order correlations retained the same direction as the standardized β coefficients, despite substantial differences between the two measures, further supports the notion that the observed effects are unlikely to be solely artifacts of the model or driven by the inclusion of many intercorrelated predictors.

Hemispheric asymmetry and its relationship with behavior have historically been discussed within a region-centered and dominance-based framework (Corballis & Häberling, 2017). Nevertheless, this study reveals lateralized multiregional associations across most behavioral measures (Fig. 1–4; Extended Data 1, 2). Moreover, it uncovers that this relationship is often directionally asymmetric. In particular, the regular occurrence of DA between contralateral homotopic regions (hDA) across an extensive set of measures is noteworthy (Fig. 1–4). This suggests that even when a classic hemispheric asymmetry, in terms of dominance, is not evident and both contralateral homotopic regions are associated with a measure, a directional asymmetry prevails: the left and right regions may show opposing associations with the same behavioral outcome. In other words, homotopic regions rarely “agree” in how their volume variation relates to behavior.

Instead, they present consistent behavioral directionality, forming push–pull pairs that extend across many behavioral tasks, from working memory and language to social cognition and fluid reasoning (Fig. 1–4, 8A–C). This may add a new dimension to the previously described aspects of hemispheric asymmetry. For instance, it might explain why substantial functional asymmetry may emerge despite minimal structural asymmetry (Perez et al., 2023; Sun et al., 2017). In addition, this suggests a new framework for asymmetry studies based on behavioral directionality rather than hemispheric dominance or competition. It is intriguing to note that the asymmetric organization of RV is not limited to homotopic regions. Indeed, an extensive ipsilateral distribution of consistent directional asymmetry between heterotopic regions (iDA) across many measures was observed. This includes, but is not limited to, instances in which two regions exhibited a multiregional polyhemispheric crossed asymmetry with dual hDA and iDA components (Fig. 1–4; Fig. 8A–C). For instance, cMFG and STG, besides contralateral homotopic asymmetry, display a consistent ipsilateral opposition with every WMRT measure, when the association reaches statistical significance (Fig. 3). In addition, as mentioned before, consistent iDA can occur independently where hDA is not present (Fig. 9).

**Fig. 8A.**
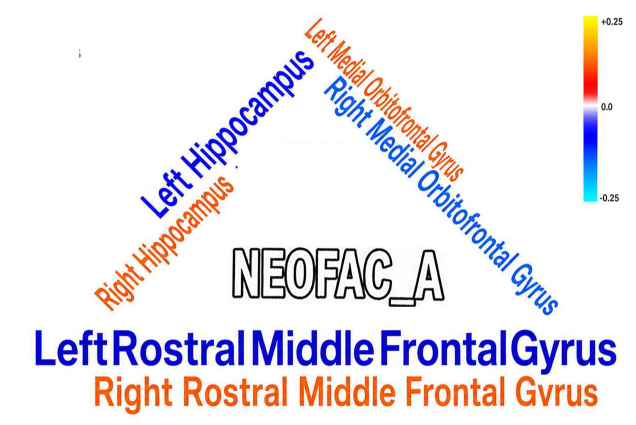
Multiregional DA. Crossed tri-regional six-hemisphere DA with dual hDA and iDA components: Rostral middle frontal and medial orbitofrontal gyri and hippocampus versus NEO Five-Factor Inventory Agreeableness (NEOFAC_A).

**Fig. 8B.**
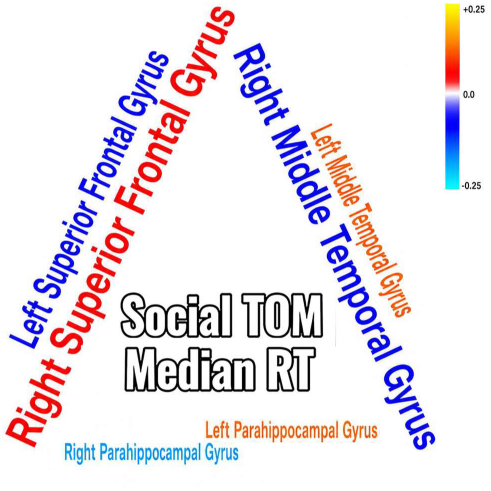
Multiregional DA. Crossed tri-regional six-hemisphere DA with dual hDA and iDA components: Middle temporal, superior frontal, and parahippocampal gyri versus Social Task Theory of Mind Median Reaction Time (Social_Task_TOM_Median_RT_TOM).

**Fig. 8C.**
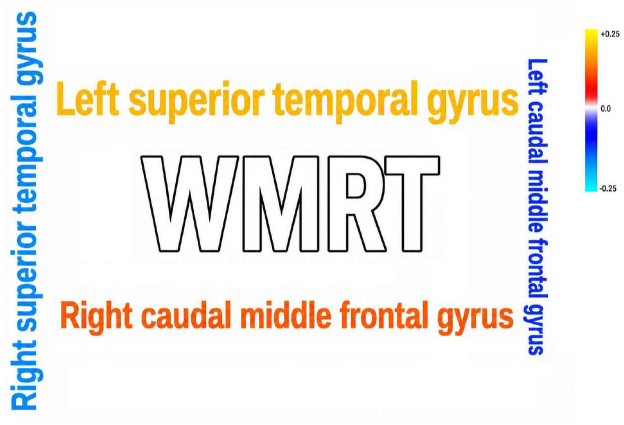
Multiregional DA. Crossed bi-regional four-hemisphere DA with dual hDA and iDA components: Caudal middle frontal and superior temporal gyri versus Total Working Memory Task Median Reaction Time (WM_Task_Median_RT).

**Fig. 9.**
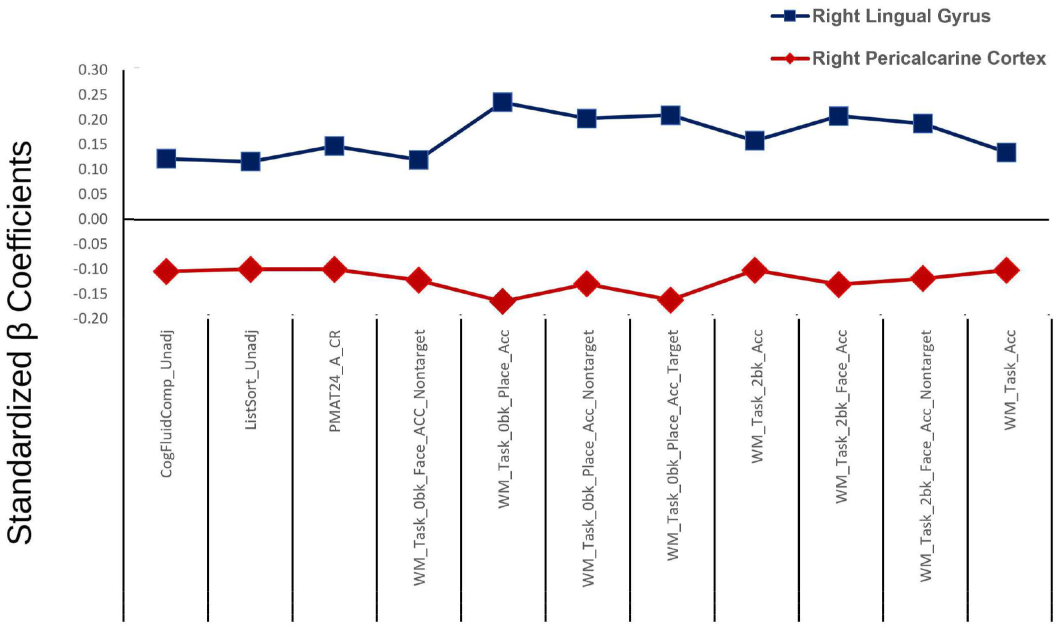
Consistent Independent iDA. Left pericalcarine cortex and left lingual gyrus exhibited iDA relative to cognitive measures.

Therefore, it appears that asymmetric organization of behavioral directionality is a pervasive biological phenomenon spanning both contralateral and ipsilateral neural networks. Thus, the asymmetric brain–behavior relationship may not be adequately captured by a simplistic, single-region, unidirectional framework. Instead, it is crucial to consider directional aspects, which may be distributed across bilateral and ipsilateral networks, thereby creating asymmetric circuits. Thus, it appears that network directional asymmetry might be a more relevant concept than regional asymmetry alone (Fig. 8A–C; Fig. 9).

In addition, it is important to consider that brain structural volume reflects cumulative developmental, genetic, and experiential influences (Lenroot & Giedd, 2008; Peper et al., 2007). Therefore, the observed consistent bilateral and/or ipsilateral structural-behavioral directionality in this study is unlikely to reflect short-term strategy differences alone. It instead may reflect a long-term specialization. Multiple pieces of evidence in this study highlight the significance of directional networks in the human brain. First, hDA is task-general, appearing across extensive brain areas and a broad range of traits ranging from emotional and personality measures to cognitive and sensorimotor parameters (Fig. 1–4). This implies that hemispheric behavioral directionality is not an epiphenomenon of specific tasks but may represent a core organizing axis of brain–behavior mapping. For example, the multiregional directional counterplay in GMV, including the cMFG–STG crossed DA association with WMRT (Fig. 3)—a key index of cognitive performance—as well as the rMFG–MOFG and HPC–MOFG associations with NEOFAC_A (Fig. 1), suggests that DA may contribute to neural mechanisms sustaining cognitive and emotional efficiency through dynamic polyhemispheric coordination across distributed brain regions. It is intriguing to note that in the cerebral cortex, hDA is mostly observed in regions that are especially developed in the human brain, i.e., the frontal and temporal cortices (Fig. 5). On the other hand, regions that display symmetric RV are comparatively uncommon and located in older cortical regions (Extended Data 2). In addition, the number of behavioral traits involved in this relationship is considerably smaller (3%). The fact that DA is particularly pronounced in brain regions that underwent disproportionate evolutionary expansion in the human lineage further suggests that this lateralization itself may represent an important neurobiological adaptation that contributes to the unique complexity, sophistication, and context-sensitive regulation of human cognition and behavior.

In fact, prior research suggests that E/I balance should be understood as a dynamic, network-level property of human brain function, relevant to cognition, development, and brain-based disorders, rather than as a feature of isolated brain regions alone (Cohen Kadosh, 2025). Consistent with this perspective, our findings raise the possibility that lateralized directional networks may contribute to the fine-tuning of neural stability, information-processing efficiency, and flexible regulation of complex human behavior. In addition, our data align with a growing body of evidence indicating that the human brain hemispheres exert opposing regulatory influences on different behavioral components including emotion, sexual behavior, arousal, consciousness, and primary physiological functions such as heart rate, blood pressure, and immune system activity (Toga & Thompson, 2003; Hugdahl, 2000; Liu et al., 2009; McAvoy et al., 2016; Joliot et al., 2016; Vallortigara, 2006; Rogers, 2014).

Moreover, a recent study utilizing resting-state fMRI data from the HCP demonstrates that distinct brain networks exhibit opposing laterality dynamics, with the left hemisphere’s language and default mode networks showing strong leftward laterality while the right hemisphere’s cingulo-opercular, dorsal attention, and visual networks display rightward laterality (Wu et al., 2022). Our data extend these findings in quantitative structural form, supporting the notion of behavioral directionality, where different areas may fine-tune cognitive and behavioral processes, facilitating adaptive responses to environmental demands.

The E/I balance is also involved in the pathophysiology of diverse neurological and psychiatric disorders when this finely tuned system becomes disrupted. Clearly, further research, particularly functional imaging studies, is required to better understand how asymmetric directional brain networks modulate human behavior. If complementary data support the hypothesis, it would be reasonable to speculate that variations in bidirectional E/I fine-tuning could contribute to differences in human psychological characteristics as well as to the pathophysiological basis of many neurological and psychiatric diseases. In fact, although numerous studies emphasize that an E/I imbalance at the cellular and synaptic levels contributes to neurodevelopmental and psychiatric disorders, a growing body of work supports the view that this imbalance must be considered at multiple scales. In fact, disruptions at the network, macro-circuit, and hemispheric levels can contribute to neurological and psychiatric symptomatology (Sohal & Rubenstein, 2019; Wang, 2020). Therefore, improving our understanding of how behavioral directional networks fine-tune behavior could open new perspectives for developing more effective therapeutic strategies for disorders that largely lack effective treatments.

## Notes

### Competing Interest Statement

The authors have declared no competing interest.

